# Genetic basis of resistance in hosts facing alternative infection strategies by a virulent bacterial pathogen

**DOI:** 10.1101/2024.08.15.607931

**Authors:** Eglantine Mathieu-Bégné, Sabrina Gattis, Dieter Ebert

## Abstract

Having alternative infection routes is thought to help parasites circumvent host resistance, provided that these routes are associated with different host resistance loci. This study examines whether alternate infection routes of the parasite *Pasteuria ramosa* are linked to distinct resistance loci in its crustacean host, *Daphnia magna*. We focus on the *P. ramosa* isolate P15, which can attach and penetrate the host through either the hindgut or the foregut. Using a global panel of 174 *D. magna* genotypes supplemented with breeding experiments, we analyzed resistance patterns for each of these infection routes. Our findings confirm our hypothesis: in *D. magna*, hindgut attachment is determined by the D locus, while foregut attachment is controlled by a newly identified G locus. We established a gene model for the G locus that indicated Mendelian segregation and epistatic interaction with at least one other resistance locus for *P. ramosa*, the C locus. Using genomic Pool-sequencing data, we localized the G locus within a known Pasteuria Resistance Complex on chromosome 4 of *D. magna*, whereas the D locus is on chromosome 7. Two candidate genes for the G locus, belonging to the Glycosyltransferase gene family, were identified. Our study sheds new light on host–parasite coevolution and enhances our understanding of how parasites evolve infection strategies.

**Author summary:** Parasites continuously evolve strategies to overcome host resistance, including the use of alternative infection routes. However, this strategy is advantageous only if host resistance loci are specific to each entry point; otherwise, a single host gene could provide resistance to all infection routes. In this study, we tested this hypothesis using the freshwater crustacean *Daphnia magna* and a strain of the parasite *Pasteuria ramosa* that can infect its host via the esophagus (foregut) or the hindgut. By conducting a phenotypic assay of *P. ramosa* attachment on a global panel of *D. magna* genotypes, we demonstrate that foregut and hindgut infections are associated with independent genetic host resistance loci. Through a breeding experiment on a subset of *D. magna* genotypes, we were able to propose a gene model for the newly discovered G locus linked to foregut attachment, while the previously identified D locus is linked to hindgut attachment. We also discovered that the foregut infection route is influenced by an epistatic interaction between the G locus and another *P. ramosa* resistance locus, the C locus. Using genomic data, we confirmed that the G and D loci are not overlapping, with the G locus being part of the Pasteuria Resistance Complex on chromosome 4, whereas the D locus is on chromosome 7. Two potential genes involved in glycosylation processes were identified as candidates for the G locus. Overall, our study confirms a key postulate in the understanding of host–parasite co-evolution, highlighting the importance of infection strategies in host resistance.

## Introduction

The process by which a parasite (incl. pathogens) infects a host is often depicted as a stepwise process starting with host encounter, followed by host entry via a specific route, then within-host growth, parasite proliferation and transmission (Fenton et al. 2012; Hall et al. 2017; Lievens et al. 2018; Hite 2020; Izhar et al. 2020). In this model, the parasite follows a linear path, where each step interacts with different aspects of its host. The host can evolve resistance by blocking the parasite at any of these different steps; the earlier this happens, the less damage the parasite causes (Lievens et al. 2018; Izhar et al. 2020). However, parasites may evolve ways to circumvent host resistance, leading to co-evolution.

Coevolution between host and parasite is based on reciprocal selection and is thought to drive diversity among organisms (Hamilton 1982; Mouritsen and Poulin 2005; Spurgin and Richardson 2010; Bever et al. 2015). Knowing the genomic architecture that underlies resistance traits in hosts and virulence traits in pathogens is thus critical to our understanding of host-parasite coevolution (Kurtz et al. 2016; Ebert and Fields 2020). It comes as no surprise that identifying the loci of host and parasite interactions has been a persistent focus in disease biology (e.g., Beraldi et al., 2007; Routtu & Ebert, 2015; Shea et al., 1996; Su et al., 2002; Turner et al., 2011). Research has documented strong and specific genetic associations between hosts and their parasites, for example, between mosquitoes and the blood parasite *Plasmodium falciparum* (Lambrechts et al., 2006), between the filter feeding crustacean *Daphnia magna* and its obligate bacteria pathogen *Pasteuria ramosa* (Carius et al. 2001), between the blood parasite *Schistosoma* and its intermediate host the aquatic snail *Biomphalaria* (Webster and Woolhouse 1998), and in human MHC in relation to diverse pathogens (Råberg 2023). Relationships between the host resistance locus and the parasite virulence locus have also been documented at the gene level, providing evidence for the gene-for-gene model and the matching allele model (Dodds et al. 2006; Luijckx et al. 2013; Råberg 2023).

The stepwise model of infection often neglects the fact that, from the parasite’s point of view, the host constitutes a heterogeneous landscape (Lydecker et al. 2019). For ectoparasites, the host’s body surface presents several potential microhabitats that are not equivalent; some locations may be preferred as an entry point by a given parasite (Loot et al. 2004; Pigeault et al. 2020). Endoparasites typically have defined entry points to the host, such as the skin, the lung, the gut epithelium, and the mucosal surfaces of genital organs (for sexually transmitted diseases). This goes down even to specific cell types a parasite can colonize (Iyer et al. 2007). Because parasites view hosts as heterogeneous, they have potentially diverse infection routes, and by evolving the ability to use alternative paths to enter the host, they can overcome host resistance. However, alternative infections routes only benefit the parasite if host resistance to one route does not simultaneously close down all other routes—i.e., if the resistance mechanism is route-specific. Therefore, when parasites successfully use different host entry routes, we expect host resistance polymorphisms to be specific to each individual route, i.e. different entry points are associated with different host genes. Thus, the host can be expected to have different genes associated with resistance to the same parasite line.

The use of alternative host entry points suggests that pathogen infection can occur as a non-linear process (with different potential branches) rather than a linear process. The evolution of host resistance and parasite virulence and its genomic basis remains quite unexplored in the context of non-linear infection models. Part of the reason is because the use of alternative infection paths by parasites is rarely considered explicitly within a non-linear model of infection. For instance, it has been suspected that oral and systemic infections of *Drosophila melanogaster* by *Pseudomonas entomophila* are associated with different resistance genes (see (Martins et al. 2013)); however, here the different infection routes are actually two different steps of a linear infection model: in one case, the encounter occurs as *in natura* (i.e., oral infection) while in the other case the pathogen circumvents the encounter step to start directly at the step of pathogen recognition by the host (i.e., systemic infection by pricking the cuticle of the host with an pathogen loaded needle). Hence, there is often confusion in the literature between a “shortened” linear infection model where the encounter is circumvented and a non-linear infection model, where a pathogen naturally infects through a different host entry point (e.g. (Eleftherianos et al. 2022). Linear and nonlinear infection models are conceptually different (see Figure 1). Here we propose that a nonlinear infection model has different evolutionary outcomes from linear infection models, which can lead to host loci specific to a pathogen’s particular infection route.

**Figure 1:**
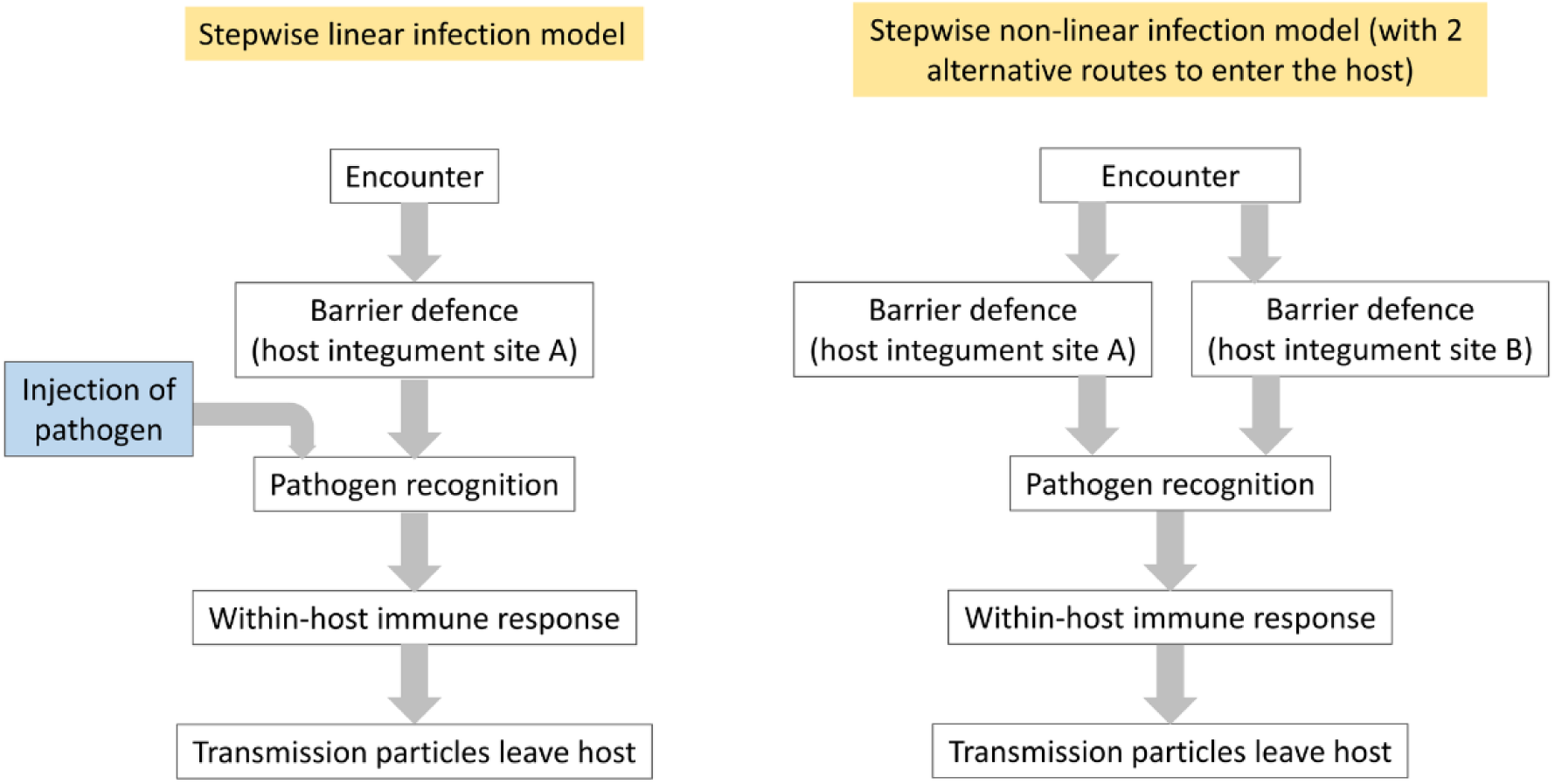
Linear (left) vs. nonlinear (right) infection models. The blue text box indicates systemic infections, e.g. by pricking the skin with a needle dipped in a pathogen culture, circumvent the first line of defense in linear infection models, shortening the model, but not presenting an alternative path for the pathogen. The non-linear infection model (right) shows two alternative routes of infection, where the pathogen can enter the host in two different sites.

The freshwater crustacean *D, magna* is often infected with the obligate bacterial parasite *P. ramosa*. According to data presented by Bento et al. (2020), a particular isolate (P15) of the parasite can enter through the host cuticle via two different natural routes: the esophagus (foregut) and the hindgut. Attachment to the gut cuticle is a strong indicator of successful infection (Duneau et al. 2011; Bento et al. 2020). Previously, resistance loci have been identified for parasites that enter the host through a specific site; for example, *P. ramosa* isolates P15 and P21 entering through the hindgut are associated with host resistance loci D and F (Bento et al. 2020; Fredericksen et al. 2023), while the A, B and C loci determine resistance to *P. ramosa* isolates C1 and C19 that enter the host exclusively through the foregut (Bento et al. 2017). Because resistance to different infection routes has only been characterized so far for different *P. ramosa* isolates it has not been possible to distinguish whether the host resistance loci were an effect of the *P. ramosa* line or of the infection route. More recently, Fredericksen et al. (2021) have discovered the potential use of multiple infection routes in about two thirds of *P. ramosa* isolates, often with different parasite isolates using different combinations of routes (out of 5). This plurality of infection routes within *P. ramosa* lines indicates that a linear infection model does not capture the complexity of the coevolutionary interactions between the host and the parasite. Instead, it is necessary to determine if parasites using different infection routes interact with the same or with different host genes.

In this study, we test—at both the phenotypic level and the genomic level—the hypothesis that different infection routes used by a single *P. ramosa* isolate are associated with different *D. magna* resistance loci. At the phenotypic level, we characterized the different infection routes (hindgut and foregut) of the *P. ramosa* isolate P15 on a worldwide panel of 174 *D. magna* genotypes. We specifically tested phenotypic associations between attachments made via different routes of infection and between three different *P. ramosa* isolates (our focal isolate P15, and two other isolates (C1 and C19) routinely used in our lab that have a known relationship to a locus often involved in epistatic interactions with other resistance loci (Metzger et al. 2016; Ameline et al. 2021)). We further conducted selfing experiments on a restricted panel of *D. magna* genotypes to investigate segregation patterns between P15 hindgut and foregut attachments. If different infection routes are indeed associated with different, unlinked host loci, we expect to see that P15 foregut and hindgut attachments are independent of each other. Furthermore, using a pool-sequencing approach, we localized the host loci responsible for P15 foregut attachment within *D. magna* reference genome. The locus connected with P15 hindgut attachment (D locus, on chromosome 7 (Bento et al. 2020)) was already known. If our hypothesis is true, we expect the locus associated with P15 foregut attachment to be in a different position. With this study, we challenge the idea that parasitic infections follow a linear, stepwise process, which is of prime interest, since parasites’ ability to escape host resistance mechanisms can raise health concern as new strategies of infection arises.

## Results & Discussion

### Characterization of the routes of infection of P15

To assess whether different infection routes are associated with different host resistance polymorphisms, we first identified co-occurrences in the attachment patterns (i.e., routes of infection) of *P. ramosa* isolates P15, C1 and C19 using a panel of 174 *D. magna* genotypes collected from the host’s Holarctic distribution (Bento et al. 2020). Using data from Bento et al. (2020) on foregut and hindgut attachments of the three *P. ramosa* isolates, we found polymorphism for all three parasite isolates, but C1 and C19 showed only foregut attachment, whereas P15 showed foregut (P15F) and hindgut (P15H) attachment to some host genotypes. Attachment by the three isolates was not independent of each other: we found a significant positive co-occurrence between C1 and C19 attachments (see Figure 2). Conversely, we found significant negative co-occurrence between P15F attachment and the three other resistotypes (C1, C19 and P15H, Figure 2). The negative co-occurrence between P15H and P15F attachments, and the fact that all four possible resistotype combinations were observed, hints that the loci responsible for P15F attachment and P15H attachment might indeed be different (i.e., the locus D). Remarkably, no susceptibility of P15F and the other foregut-attaching isolates, C1 or C19, was observed (Fig. 2),

**Figure 2:**
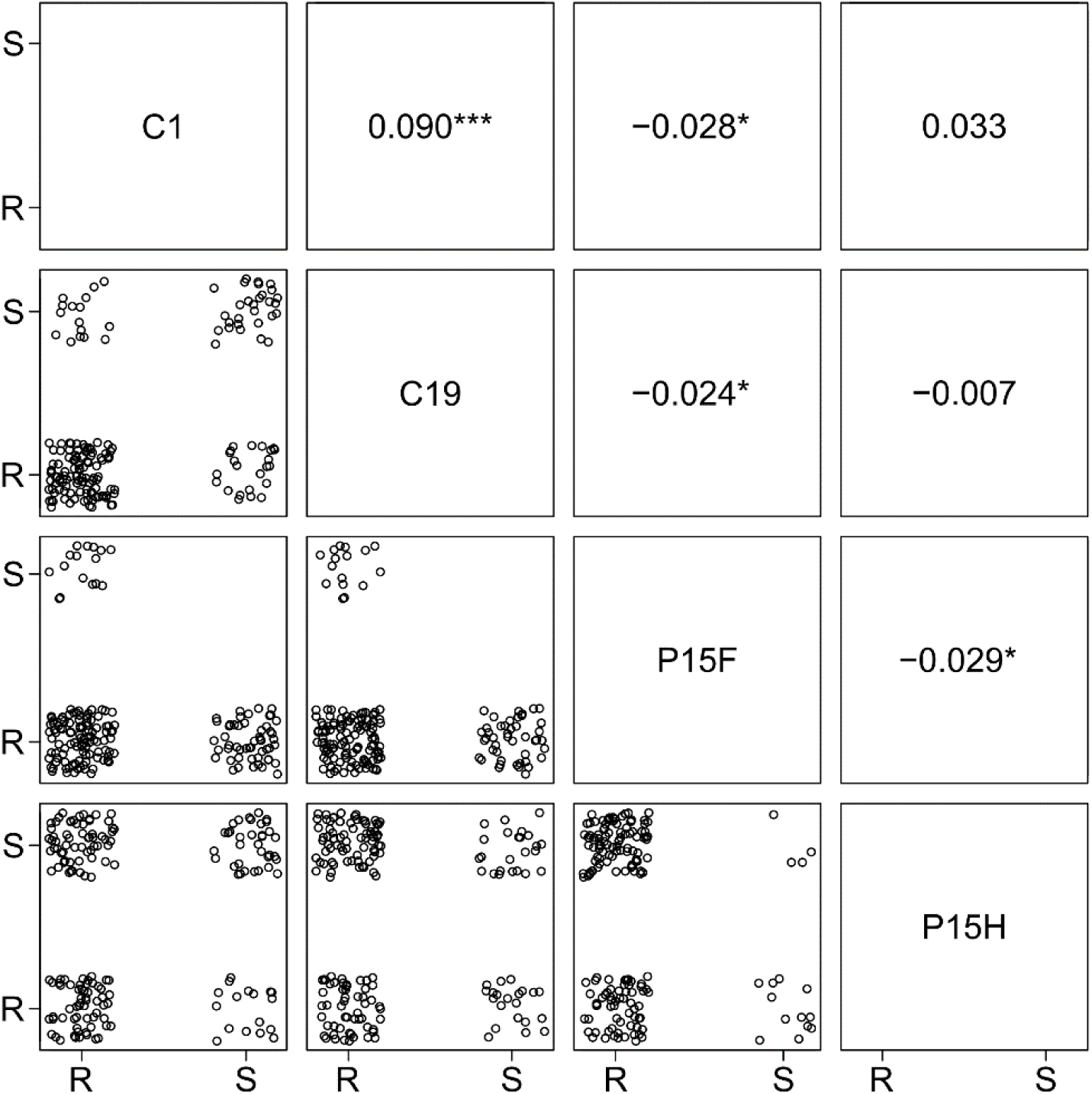
Co-occurrences between the attachments of C1, C19, P15 hindgut (P15H) and P15 foregut (P15F). Dots indicate a *D. magna genotype* as either resistant (R) or susceptible (S) to each *P. ramosa* isolate. Numbers on the upper part of the plot are standardized effect size from the co-occurrence test (i.e., the differences between expected and observed frequency of co-occurrence). Positive effect size indicates a positive co-occurrence, and negative, a negative co-occurrence. Stars indicate the test’s level of significance, if any (*** is <0.001, ** is <0.01 and * is <0.05). P-values are provided in Table S1.

To understand the segregation of resistance loci and test the segregation between P15F and P15H attachment, we conducted a breeding experiment on 39 *D. magna* genotypes. Selfed F1 offspring were produced by allowing females to mate with their asexually produced sons or brothers (Table S2). Selfing of all 39 clones produced viable selfed F1 offspring in 31 clones (Table S3). The reassessment of parents and offspring for P15, C1 and C19 attachment confirmed previously reported resistotypes, with C1 and C19 showing only foregut attachment, while P15 showed foregut and hindgut attachments. Selfed F1 offspring of four parental genotypes segregated for P15F attachment (i.e., TN-RA-2, TN-RA-21, ES-HT-1 and CY-PA3-3, see Table 1 and Table S3). While TN-RA-2 and TN-RA-21 were resistant to P15F, ES-HT-1 and CY-PA3-2 were susceptible to P15S. Their C1/C19 resistotypes were RR in all cases. These 4 clones showed no segregation for P15 hindgut attachment, even though they differed, with TN-RA-2, TN-RA-21 and ES-HT-1 being P15H resistant (P15H-R), whereas CY-PA3-2 was susceptible (P15H-S)(Table S3). This suggests that the D locus in these clones is homozygote (dd in case of P15H-R, and DD in case of P15H-S) and confirms that the putative locus for P15 foregut attachment segregates independently of the D locus.

**Table 1:** Parental clone and selfed-offspring resistance phenotypes (i.e., resistotypes) for the four *D. magna* clones among the selfed offspring that segregated for *P. ramosa* P15F attachment. Hypothesized genotypes are given according to genetic model 1 and 2 for the putative G locus. Finally, the percentage of the observed and expected P15 foregut attachments based on the gene model 1 or gene model 2 for the G locus are provided. For the full table of the 39 genotypes used in the breeding experiment, refer to the Table S3.

| Parent clone ID | Resistotype |  | Hypothesized genotype (CG loci, model 1 or model 2) | Selfed offspring (F1) resistotype |  | Nobs | %obs |  | %exp | Hypothesized genotype (CG loci, model 1 or model 2) |
| --- | --- | --- | --- | --- | --- | --- | --- | --- | --- | --- |
|  | C1, C19 | P15F |  | C1, C19 | P15F |  |  |  |  |  |
| CY-PA3-2 | RR | S | Ccgg or CcGG | RR | S | 10 | 71.43 |  | 75 | C-gg or C-GG |
|  |  |  |  | SR | R | 4 | 28.57 |  | 25 | ccgg or ccGG |
| ES-HT-1 | RR | S | Ccgg or CcGG | RR | S | 24 | 63.16 |  | 75 | C-gg or C-GG |
|  |  |  |  | SR | R | 14 | 36.84 |  | 25 | ccgg or ccGG |
| TN-RA-02 | RR | R | CCGg | RR | R | 27 | 87.1 |  | 75 or 25 | CCG- or CCgg |
|  |  |  |  | RR | S | 4 | 12.9 |  | 25 or 75 | CCgg or CCG- |
| TN-RA-21 | RR | R | CCGg | RR | R | 22 | 81.48 |  | 75 or 25 | CCG- or CCgg |
|  |  |  |  | RR | S | 5 | 18.52 |  | 25 or 75 | CCgg or CCG- |

### Role of epistatic interaction in P15F attachment

As mentioned above (Figure 2), we did not find P15F susceptibility together with C1 or C19 susceptibility. The absence of *D. magna* genotypes susceptible to C1, C19 and P15F suggests a possible epistatic interaction between the loci responsible for C1 and C19 resistance and the locus responsible for P15F resistance. The loci responsible for C1 and C19 susceptibility are the A, B and C loci, located within a cluster on chromosome 4 (known as the Pasteuria Resistance Complex, PRC, earlier referred to as the ABC supergene (Bento et al. 2017; Fredericksen et al. 2023)).

Combining C1 and C19 resistotypes into double types (RR, SS, SR and RS), we observe that SS, SR and RS never coincide with P15F-S, while RR does. Bento et al (2017) showed that the C locus segregates exactly in this way with the phenotypes, such that RR resistotypes are *CC* or *Cc* (C allele is dominant), while SS, RS and SR resistotypes are always *cc*. This observed pattern suggests that the C locus genotype *cc* is resistant to P15F. Since the C locus does not otherwise influence P15F resistance (*CC* and *Cc* genotypes can be P15F-S or P15F-R), we conclude that another locus must influence P15F resistance, but that it acts epistatically with the C locus. We hypothesize that the *cc* genotype nullifies the effect of the P15F resistance locus, regardless of the genotype at P15F resistance locus.

Note that genotypes susceptible to C1 or C19 and P15F at the same time was never observed in Bento et al. (2020) data that we are analyzing here and this was also almost unanimously the case for any genotype tested routinely in our lab hence largely confirming the epistatic interaction between the C and the G locus. However, it is worth noting that few exceptions representing a bit less than 1% of the genotype tested in our lab (i.e., three genotypes) were observed where some susceptible genotypes to P15F were also susceptible to C1 (but not to C19). This very rare exceptions may be explained by a epistatic interaction involving the B locus that appear rare due to the silencing effect of the C and the A locus on the B locus (see Bento et al. 2017 for details on the model) but this would require a dedicated dataset to be fully explored. In any case, these rare exceptions do not invalidate the hypothesis of an epistatic interaction between the C and the G locus.

Among the 39 *D. magna* genotypes used in the breeding experiment, we also confirmed that P15F attachments (=P15F) were seen only among clones with RR resistotypes at C1/C19 (12 of 27 RR clones), while clones with C1/C19 resistotypes RS, SR, and SS (genotype *cc*) were always resistant to P15F attachment (12 of 39 clones, Table S3).

Overall, these results suggest that the genetic architecture of P15F attachment is influenced by two loci, with the C locus epistatically masking the effect of the other locus when an individual carries the *cc* genotype. We will hereafter refer to the new locus associated with P15F attachment as the G locus (following the naming sequence that labeled the last discovered locus the F locus).

### Segregation patterns and gene model for P15F resistance locus

We used the breeding experiments, to further investigate segregation patterns and potential dominance of the locus responsible for P15F attachment. As described above, four genotypes showed segregation for P15 foregut attachment in their selfed offspring. The proportions of susceptible and resistant self F1 offspring to P15F attachment were close to Mendelian, suggesting that a single locus with two alleles and complete dominance could be responsible for P15F resistance (Table 1). Mendelian segregation is commonly observed on the currently-known resistance loci of *D. magna* to *P. ramosa* (Bento et al. 2017; Bento et al. 2020; Ameline et al. 2021; Fredericksen et al. 2023). Two parental clones that segregated for P15 foregut attachment in their selfed offspring were susceptible (CY-PA3-2 and ES-HT-1, Table 1) while two others were resistant (TN-RA-2 and TN-RA-21 clones, Table 1). The epistatic interaction between the C locus and the G locus deduced above could explain this pattern. The segregation for P15 foregut attachment in CY-PA3-2 and ES-HT-1 is compatible with the masking effect of the C locus on the G locus. Thus, P15F segregation is observed because of segregation of a heterozygote *Cc* parental genotype, while the G locus is homozygote. The TN-RA-21 and TN-RA-2 clones showed no segregation at the C locus, with parents and all offspring being resistotype RR (Table 1). This suggests that these two clones are homozygote for the dominant allele of the C locus (genotype *CC*) and heterozygote for the G locus. The G locus would therefore be segregating in the offspring of TN-RA-21 and TN-RA-2 (genotype Gg) (Table 1). While the ratios between resistant and susceptible offspring in TN-RA-21 and TN-RA-2 suggest that the locus responsible for P15F attachment is dominant for resistance, the overall low number of offspring segregating for P15F attachment and the strong selection against inbreeding (all selfed F1 are 50 % inbreed!) suggest we take this hypothesis with caution.

Two alternative models for the G locus and a summary of the above deductions are presented in Figure 3. Hypothesized genotypes are reported in Table 1. The G locus is predicted to be a two allele Mendelian locus with total dominance (like other *D. magna* resistance loci against *P. ramosa*); the models differ only in that resistance is dominant in model 1 while susceptibility is dominant in model 2. In both model 1 and 2, the clones TN-RA-2 and TN-RA-21 are predicted to be heterozygotes for the G locus. Likewise, epistatic interactions with the C locus are present, such that *cc* genotypes nullify the effect of the G locus. Note, the C locus has previously been shown to interact epistatically with other loci. The dominant C allele nullifies the A and the B loci, while the genotype *cc* nullifies the E locus (Bento et al. 2017; Ameline et al. 2021). The pivotal role of the epistatic interaction involving the C locus in determining P15F attachment is hence very much in line with observations about other resistance loci against *P. ramosa* in *D. magna*.

**Figure 3.**
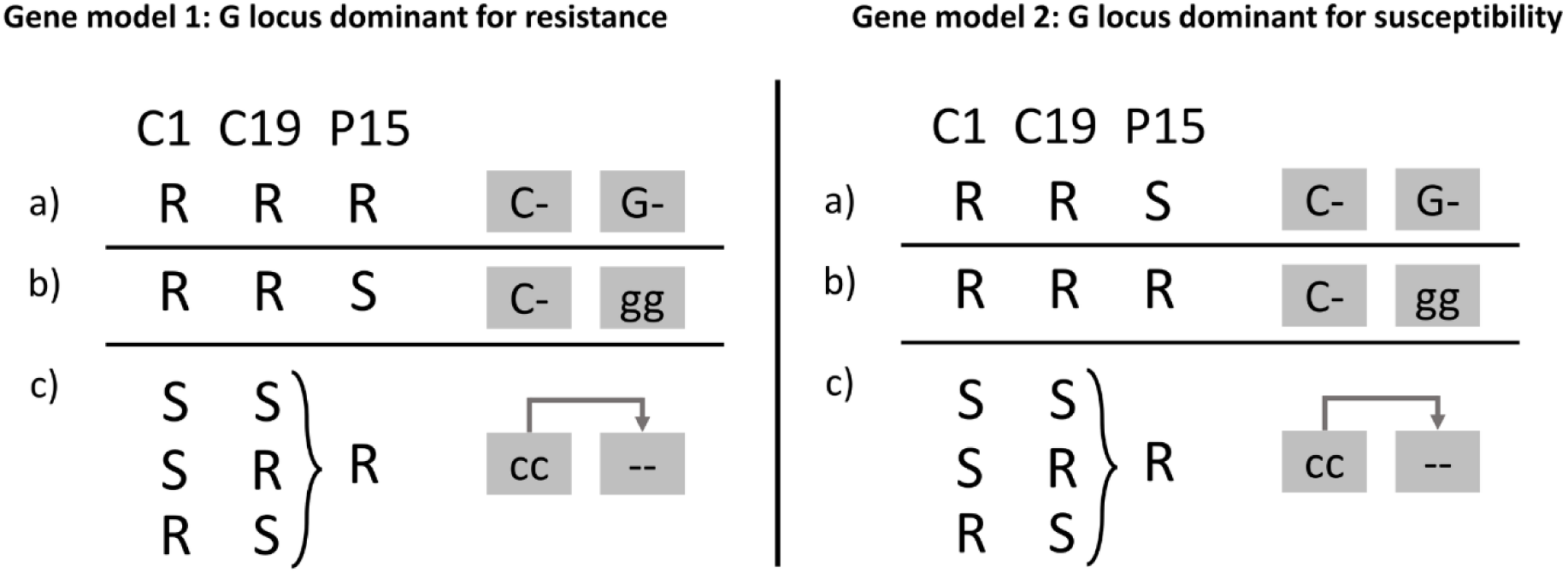
Two alternative genetic models for the G locus. Resistance to *P. ramosa* P15 foregut attachment to be determined by the G locus and its interaction with the C locus. Genotypes at these loci are represented by boxes with grey background. Each locus has two alleles, with upper case denoting dominance. The dash (-) indicates a wildcard, i.e. that the alleles on these positions do not play a role in determining the host’s resistotype. a) P15 foregut resistance is dominant (left, model 1) or susceptibility is dominant (right, model 2); b) P15 foregut susceptibility (model 1) or resistance (model 2) is conferred by the recessive alleles on the G locus. c) Epistatic interaction (dark grey arrow): C locus genotype *cc*, nullifies the effect of the G locus, such that homozygosity of the recessive alleles on the C locus masks the expression of alleles on the G locus, making the host resistant to P15 foregut attachment (true for both model 1 and model 2). The letter R and S stand for resistance and susceptibility to the *P. ramosa* isolates listed in the top row.

### Genomic Mapping of the G locus

To locate the G locus, we relied on pool sequencing of resistant and susceptible selfed offspring of clone TN-RA-21 (i.e., predicted heterozygote at the G locus, Table 1). Specifically, we used Fisher Exact Tests to detect significant changes in allele frequencies between a pool of 33 genotypes susceptible to P15F attachment and 23 genotypes resistant to P15F attachment. A clear peak of allelic frequency change was observed on contig 11 of chromosome 4 (Fig. 4), indicating that the G locus, is definitely in a different region than the D locus (chromosome 7) that is responsible for the polymorphism of P15 hindgut attachment (Fig. 4). Both phenotypic and genotypic evidence, thus, establish our prediction that alternative paths of infection can be associated with different host loci.

**Figure 4:**
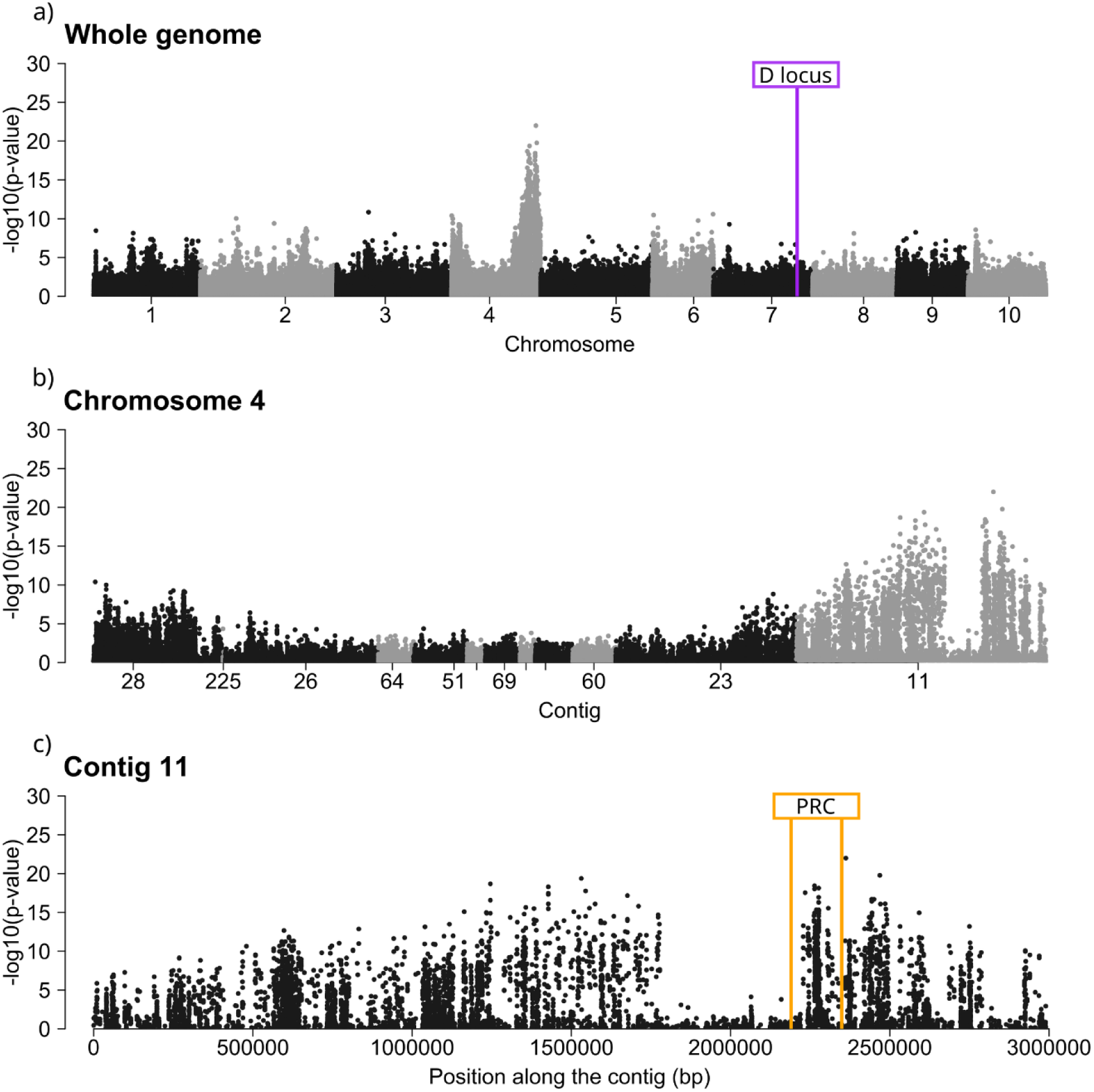
Manhattan plots representing the -log10(p-values) of the Fisher Exact Tests conducted on alleles frequencies of the resistant and susceptible pools (replicate 1, see Fig. S1 for all three replicates) as a function of genomic positions across the whole genome and (a) zoomed in to chromosome 4 (b) and to contig 11 on chromosome 4 (c). The position of the D locus with the A,B,C, F locus cluster (i.e., Pasteuria Ramosa Complex, PRC) also indicated.

A broad peak spanning about 3 MB is visible in a close-up of contig 11 on chromosome 4 (Fig. 4B). An area about 400 kb wide within the peak region shows allele differentiation close to zero (called hereafter the gap region, Fig. 4B and C). Mapping depth and SNP quality were similar to the flanking regions, but the number of SNPs was strongly reduced in the gap region, suggesting that the TN-RA-21 genotype is homozygote in the gap region (Fig. S3).

### Refining the position of the G locus

Pool-sequencing data are often used to compute differentiation metrics (such as Fisher Exact Test or *F*st) that allow us to associate SNPs with phenotypic variation across different pools (Schlötterer et al. 2014). Although these differentiation metrics usually provide insight about the location of a specific locus, mapping accuracy is proportional to the number of recombination events that occurred between the two pools. In our case, because we used the self-offspring of one clone, the number of recombination was limited, so indeed, the peak we detected on contig 11 of chromosome 4 was very broad (Fig. 4). To refine the position of the locus, thus, we developed a method using contrasting assumptions from the two hypothesized gene models for the G locus (Fig. 5). inspired by a technique that has been used to map sex determinant loci, which are typically single Mendelian loci with perfect dominance (e.g. Wen et al. 2022). Since the G locus is predicted to be a Mendelian segregating locus, we expected one pool to be homozygote at the G locus (genotype *gg*) and the other to be a mix of heterozygotes (*Gg*) and homozygote (*GG*). Using this reasoning, we computed private allele content at the contig 11 in each pool, where one pool should contain only the private G allele (the pool being *GG* and *Gg*), which should be absent in the other pool (*gg*). Private alleles were found in the susceptible pool (∼2260000-2290000 bp on the contig 11, Fig. 5), but not in the resistant pool, suggesting that the G locus is dominant for susceptibility, which supports genetic model 2. This fine mapping of the G locus revealed its position within the PRC (Fig. 4) in a window of about 30 kb. The same conclusion was reached using filters applied directly to allele frequencies to select SNPs for which one pool is homozygote (major allele frequency of one pool over 90 %, see Fig S4): When keeping only homozygote variants in the susceptible pool, no other allele frequency differences were detectable. Thus, we are confident that the G locus is most likely a single locus, dominant for susceptibility (model 2 in Figure 2).

**Figure 5:**
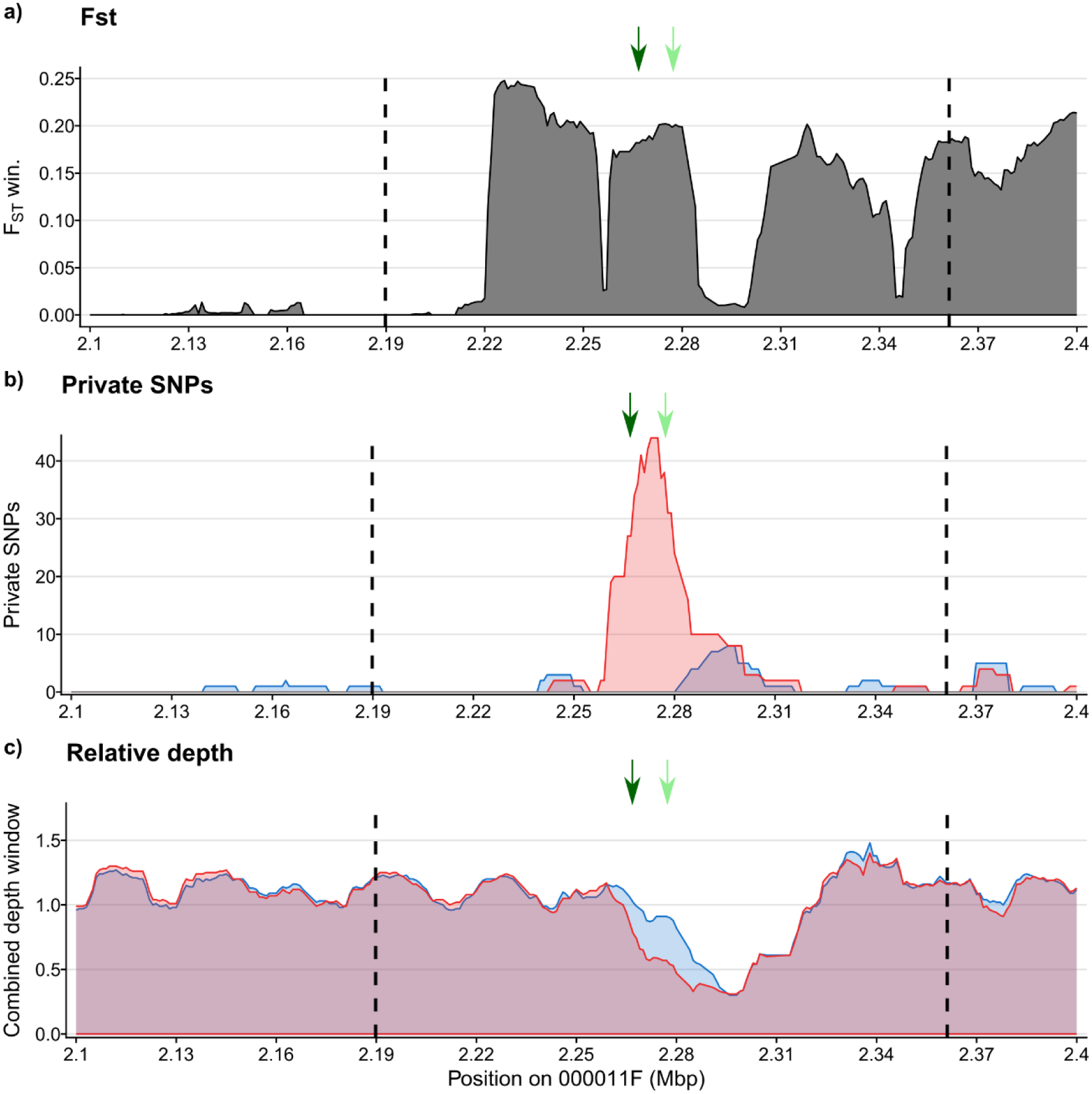
a) Fst, b) private alleles content and c) coverage metrics in the resistant and susceptible pools (blue and red respectively) along the region of the Pasteuria Resistance Complex (PRC), a cluster of several *P. ramosa* resistance loci including the A,B,C and F loci in *D. magna*. The PRC is positioned within an hyperdivergent haplotype on contig 11 of chromosome 4 and is here delineated with dashed lines. Arrows indicate the position of the two candidate genes for the G locus (dark green for the *lactosylceramide 4-alpha-galactosyltransferase-like* gene and light green for the *putative Alpha--fucosyltransferase C* gene respectively). Each graph is based on 10 kb windows with a 1 kb resolution.

The method we present here could be strengthened using coverage metrics, which also contain information about the heterozygosity of an individual at a genomic region of interest. However, since the PRC is positioned within a large, highly variable and non-recombining region (HDH-4-1, Naser-Khdour et al. submitted) that maps poorly to the reference genome, coverage metrics were not able to reliably refine the G locus location in our case. Even so, it pointed at the same position inside the PRC. If the G locus had not fallen into a Hyper Divergent Haplotype (i.e., where significant structural variation occurs between genotypes), it is reasonable to assume that we would have been able to further refine its location window. We hence propose that applying private allele and coverage differences to pool-sequences can be an efficient way to fine map loci for plausible gene models.

Interestingly, our findings suggest that susceptibility is dominant in the G locus, which brings up a commonality regarding the genetic interactions of the C locus. Susceptibility is also dominant in the E locus, while for the A and B loci (as well as the C locus), resistance is dominant. The C locus nullifies the effect of the E and the G loci when a *D. magna* clone carries two recessive alleles (*cc*) at the C locus, but the effects of the A and the B locus are cancelled when an individual carries at least one dominant allele (*CC* or *Cc*) at the C locus. Hence, we can hypothesize that the genotype *cc* nullifies the effect of loci where susceptibility is dominant, while the genotype *C-*nullifies loci where resistance is dominant. This observation may shed light on the complex epistatic interactions of *Daphnia* resistance genes against *Pasteuria*. Our results notably strengthen the idea that epistatic interactions between Mendelian segregating loci are common principle in *D. magna* resistance mechanisms to various isolates of *P. ramosa* (Metzger et al. 2016; Ameline et al. 2021).

### Candidates genes for the G locus

Overall, we recorded 24 significant SNPs whose resistant pool was almost homozygote (major allele frequency over 90 %) (Table 2, Fig. S4). Our analysis contrasts a genome pool where we expect the recessive allele (*g*) to be fixed, with another pool where homozygotes (*GG*) and heterozygotes (*Gg*) are mixed. Assuming Hardy-Weinberg proportions, the frequency of the dominant *G* allele in the other pool should be around 66 % (1/3 *GG* and 2/3 *Gg*); however, our data on the most significant SNPs suggest that its frequency is mostly slightly under 50 % (range 30 to 50 % for the private allele frequency in the susceptible pool; Table 2), making it slightly less prevalent than expected. This deviation from Hardy-Weinberg expectations might indicate that the *G* allele is at a disadvantage and may explain the overall low prevalence of P15F susceptible genotypes (see Table S1) since they are all characterized by the presence of at least one *G* allele.

**Table 2:**
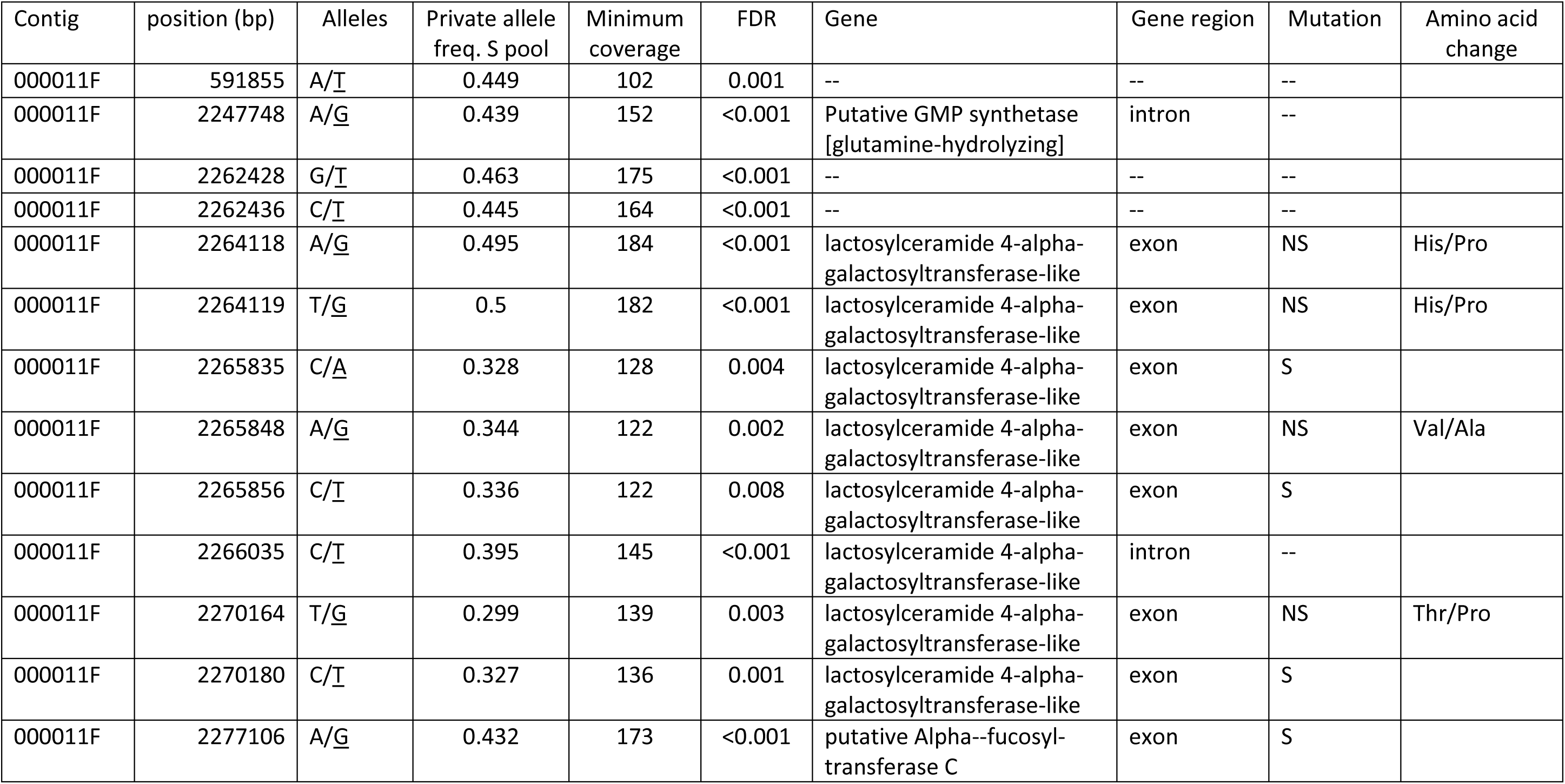
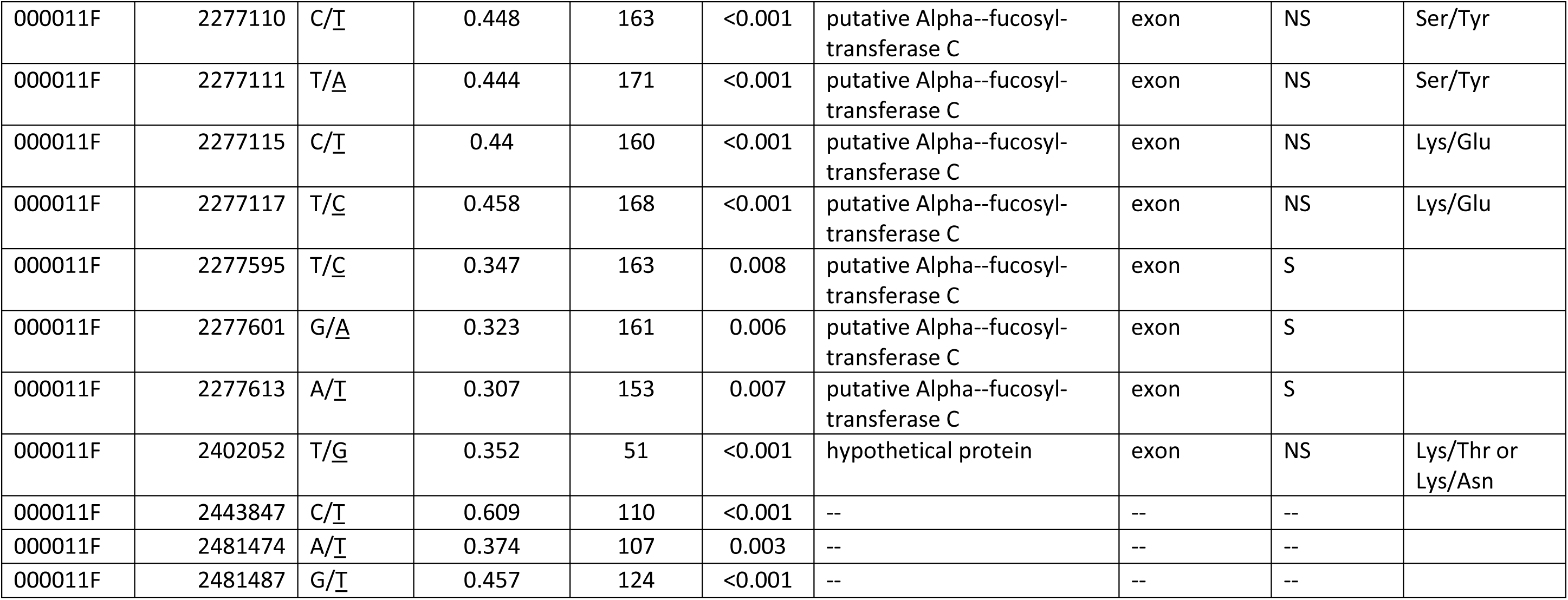
Candidate SNPs significant for allele frequency change between the resistant and the susceptible pools and for being homozygotes in the resistant pool, the contig where they are positioned, their position within the contig, the two allele states with the private allele underlined, the private allele frequency in the susceptible pool (Private allele freq. S pool), the minimum coverage at the SNP position, the corrected p-value (i.e., False Discovery Rate, FDR) of the Fisher Exact Test, the annotation of the gene if the SNP fell into a gene, the gene region when relevant (intron or exon), the type of mutation caused by the SNP (S for synonymous and NS for non-synonymous) and the amino acid change for each state of the allele when the mutation is non-synonymous.

The 24 significant SNPs fell into four different genes in the reference genome and caused non-synonymous mutations in exons in three of them (a *lactosylceramide 4-alpha-galactosyltransferase-like* (= LGT), a *putative Alpha--fucosyltransferase C* (=Fut), and a hypothetical protein, see Table 2). Most of the SNPs were concentrated in the glycosylation genes (LGT and Fut), with 8 SNPs falling into each of the two, making them strong candidates for the G locus. Some of these amino acid changes could impact protein properties, since in some cases, the alternative amino acid belongs to a different structure group than the reference amino acid. This is true for the lysine and the glutamic acids, for example, in the putative Fut sequence that belong to the basic and the acidic R group, respectively (Table 2).

The top two candidates for the G locus are relevant not only because of functional changes that might exist between resistant and susceptible genotypes to P15F attachment, but also for biological reasons. Galactosyltransferases and fucosyltransferases, are enzymes belonging to the Glycolsyltransferase family that assemble polysaccharide chains by either adding a sugar on a protein or a lipid, or by extending an existing saccharide chain (Almeida et al. 1997; De Vries et al. 2001; Ma et al. 2006; Li et al. 2018; Rini et al. 2022). These enzymes are promising candidates for resistance polymorphism, as some pathogens attach to sugars on the host epithelium before penetrating. Thus, specific sugars may be the frontline of the host’s interaction with the pathogen (Audfray et al. 2013). Furthermore, glycosyltransferase enzymes are known to have specificity for donors and/or receptors, which would provide a mechanism for a matching-allele model of co-evolution, such as *D. magna* and *P. ramosa* (Luijckx et al. 2013; Rini et al. 2022). Fucosyltransferases have already been suggested as a candidate for two other resistance loci against *P. ramosa* in *D. magna* (the C locus and the F locus, (Bento et al. 2017; Fredericksen et al. 2023)). On the pathogen side, it has been shown that collagen-like proteins such as Pcl7 in *P. ramosa* are likely involved in the attachment process (Huessy et al. 2024) and that the glycosylation of collagen-like protein in the parasite and/or of cuticle receptors of the host may explain the specificity of *P. ramosa* attachment patterns. A recent study further highlighted the high diversity in fucosyltransferase genes in *D. magna*, suggesting they might play an essential role (Naser-Khdour et al, submitted) in *P. ramosa* resistance. However, it is challenging to relate the top two candidate genes specifically to the G locus because it is within the PRC, which exists in a large non-recombinant region of the *D. magna* genome (Bento et al. 2017). The lack of homology between different haplotypes for this region poses mapping challenges that can impact our understanding of the gene’s content among different *D. magna* haplotypes. Consequently, SNPs may be preferentially detected in more conserved regions of the PRC, potentially lacking functional association with a specific target locus. Alternatively, essential genes that confer resistance to pathogens undergo strong selection pressure to be maintained at the population level. The resistance genes in the PRC have been shown to be under long-term balancing selection (Luijckx et al. 2013; Bourgeois et al. 2021; Cornetti et al. 2024). We hence suggest that the *lactosylceramide 4-alpha-galactosyltransferase-like* gene and the *putative Alpha-fucosyltransferase C* gene we identified as candidate genes for the G locus are very likely to play a role in *P. ramosa* resistance. Nevertheless, based on our data, we cannot definitively determine whether one of our top candidate genes is the G locus or not. Future research that considers multiple *D. magna* genomes, such as a pangenome approach (Sherman and Salzberg 2020), rather than a single reference, might be necessary to advance our understanding of the genomic basis of *D. magna* resistance to *P. ramosa*.

## Material and Methods

### Attachment patterns investigation on a worldwide panel

An attachment assessment of *P. ramosa* isolates P15, C1 and C19 on a worldwide panel of 174 *D. magna* genotypes was published by Bento et al. (2020). The dataset includes attachment scores (as binary scores) for each infection route used by each *P. ramosa* isolate on each *D. magna* genotype. We used the R package *cooccur* (Griffith et al. 2016) to test for positive and negative associations between attachments of C1, C19, P15 foregut (P15F) and P15 hindgut (P15H). P-values of the co-occurrence test were corrected for multiple comparison using Benjamini & Hochberg false discovery rate (Benjamini and Hochberg 1995). Results were presented as a co-occurrence matrix with effect size of the co-occurrence tests and levels of significance. Other metrics used and generated by the test, such as observed and expected occurrences, probability of co-occurrence and corrected p-values, are provided in Table S1.

### Segregation patterns investigation

Breeding experiments were conducted on a representative subset of 39 *D. magna* clones (see Table S3 for a list). These clones were kept under conditions that promoted offspring production by selfing (Ebert 2022; Santos and Ebert 2023). Attachments of C1, C19 and P15 were tested with the attachment test (Duneau et al. 2011).

Genotypes were first deduced based on the genetic model for C1 and C19 resistance (Bento et al. 2017; Ameline et al. 2021). Genotypes were then deduced from the hypothesized gene models for the new locus (G locus) responsible for P15 foregut attachment for parents and selfed offspring. We then compared phenotype frequencies as expected based on the hypothesized gene models to the observed phenotype frequencies. Recombination rates were assumed to be 0.5.

### Pool-sequencing

We conducted pool-sequencing with selfed offspring from a clone that was predicted to be heterozygote for the locus responsible for P15F attachment but homozygote for other known resistance loci (Clone TN-RA-21 from Tunesia, see Table 1), TN-RA-21 was propagated and allowed to produce resting eggs (sexual eggs = ephippia) by selfing (Santos and Ebert 2023). Resting stages were diapaused and hatched following the method of Santos & Ebert (2023). Each hatchling (F1) was cloned; then six individuals per clone were exposed to P15 spores to test for P15 foregut attachment (Duneau et al. 2011). If over 50% of the individuals showed P15 foregut attachments, they were defined as susceptible; otherwise, they were labeled resistant. Overall, 23 F1 TN-RA-21 were produced and scored as resistant, and 33 F1 TN-RA-21 were produced and scored as susceptible.

#### DNA extraction and pool-sequencing

Prior to DNA extraction, *Daphnia* were prepared following the protocol described in (Dexter et al. 2023). Three replicates of resistant and susceptible pools were produced. In order to keep the DNA contribution of each clone balanced in the pools, two individuals per clone were added to the pools as described in Table S4. DNA was extracted using the Qiagen GenePure DNA Isolation kit and following the DNA-extraction of *Daphnia* and symbionts protocol (dx.doi.org/10.17504/protocols.io.5jyl82n96l2w/v1). Briefly, whole *Daphnia* were denatured using a pestle in lysis solution. Proteinase K was then added to degrade host cuticle, and samples were first incubated overnight at 55 °C and then again for half an hour at 37 °C with RNase. They were then treated with a protein precipitation solution, and centrifuged to remove precipitates. The supernatant was treated with isopropanol and glycogen as a DNA carrier. The DNA pellet was then washed with 70 % ethanol and resuspended in hydration solution prior to DNA quantification. Libraries were prepared and sequenced as paired-end short reads (150 bp) at the Quantitative Genomics Facility service platform Basel (D-BSSE, ETH), Switzerland, using Illumina technology on a novaseq6000 sequencer.

#### Bioinformatic analysis

Paired-end raw reads were quality filtered and trimmed using trimmomatic (Version 0.39). Reads were subsequently mapped onto the *D. magna* reference genome (version 3.1) using the mapper BWA (function bwa-mem2 for speed efficiency (Li 2013; Vasimuddin et al. 2019)) with the quality filter -*q* set at 20, which allowed for the removal of ambiguously mapped reads. Bam files were sorted by coordinates and indexed. We used the software *Popoolation2* (Kofler et al. 2011) to compute allele frequencies and genomic differentiation between resistant and susceptible pools. Specifically, we created a synchronized file for resistant and susceptible pools using SAMtools mpileup function followed by the executable mpileup2sync.jar in *Popoolation2.* This synchronized file contains the allele count for all bases in the reference genome and for all pools being analyzed. We then computed allele frequencies using a minimum count of six, a minimum coverage of 20, a maximum coverage of 200 and set the pool size according to the pool size of each F1 clones (i.e., TNRA21 F1 resistant=23, TNRA21 F1 susceptible=33). We further tested for significant differences in allele frequencies using a Fisher Exact Test with geometric mean as a window summary method. The same analyses were run for each replicate of the resistant and susceptible pool to assess result reproducibility.

We tested for allele frequency differences using Fisher Exact Test first, because it is a very common metric for differentiation when working with pool-sequencing data. However, this metric resulted in a very broad peak for the location of the locus along the *D. magna* genome which was informative only at the contig level (i.e., several MBs length). As the clear peak of allelic frequency change on contig 11 of chromosome 4 was observed consistently in all three replicates (Fig. S5), we present results only from the first replicate for subsequent analyses.

To refine the position of the locus, we then used the gene models hypothesized for the locus responsible for P15F attachment. G locus model 1 and model 2 are both based on 1 locus with 2 alleles and perfect dominance. Because this model often applies to sex-determining loci, we used the PSASS (Pooled Sequencing Analyses for Sex Signal) tool that is usually used to localize sex determining loci, https://github.com/SexGenomicsToolkit/PSASS (Feron 2020). Using sliding windows across the region of interest, we visualized common differentiation statistics (Fst) and other metrics such as private alleles content in each pool (that is derived from alleles frequencies in each pool) and coverage. Using private alleles content and coverage in each pool, it is possible to refine the position of a locus for which the expected frequencies in the two pools that are contrasted are known, which in our case was provided by the gene models. In order to match Mendelian expected frequencies, we defined heterozygote frequency between 0.30 and 0.70 and homozygote frequencies as superior to 0.90. The results were visualized using modified versions of the functions present in the associate PSASS R package (psass-vis) so as to make them compatible with our own data and the R package Circlize (Gu et al. 2022). We further directly filtered SNPs based on expected major and minor allele frequencies in each pool and extracted from Popoolation2 results to confirm the results provided by PSASS.

#### Candidate genes investigation

To identify potential candidate genes for the host locus responsible for P15 foregut attachment, we first filtered SNPs based on allele frequencies expected in each pool as described above. We selected the most significant SNPs that showed a change in allele frequency between the resistant and the susceptible pool based on the corrected p-value (Benjamini – Hochberg correction) of the Fisher Exact Test at a threshold of 1%. These SNPs were then mapped onto the *D. magna* reference genome and gene model and their position visualized using the Integrative Genome Viewer (Robinson 2011). When an SNP fell onto a gene, the annotation of the gene was further checked for more up-to-date annotations using a blastx query on the NCBI online tool and with the experimental clustered non-redundant database. SNPs were categorized as falling into a non-coding region or into a coding region. When falling into a coding region, SNPs were further categorized as synonymous or non-synonymous. Note that to do so, we assumed that beside the SNPs that differed between the two pools, the rest of the sequence remained unchanged compared to the reference. As expected, the resistant haplotypes were identical to the sequences found in the reference, which is a genotype that is also resistant to P15F attachment.

## Acknowledgements

We thank members of the Ebert group for feedback on the study and the manuscript. We thank Suzanne Zweizig for language editing. This work was supported by the Swiss National Science Foundation grant numbers 310030_188887 and 310030_219529 to D.E.

## Author Contributions

Conceptualization: EMB, SG, DE

Methodology: EMB, SG, DE

Investigation: EMB, SG

Visualization: EMB

Funding acquisition: DE

Project administration: DE

Supervision: DE

Writing – original draft: EMB

Writing – review & editing: EMB, SG, DE

## Data availability

The genomic data generated in this study have been deposited in the NCBI database under accession code BioProjectID PRJNA1143412.

Scripts and data required for replicating our results are available in the Figshare repository currently accessible at https://figshare.com/s/6bed9f81f03e68201144 and will be released upon manuscript acceptance.

## Supplementary Material Caption

**Figure S1:** Read coverage calculated in 50 kb sliding windows and across contig 11 for a) the resistant pool and b) the susceptible pool. The black lines indicate the delineations of the gap region, while the orange lines indicate the position of the Pasteuria Resistance Complex.

**Figure S2:** SNP calling quality calculated in 500 SNPs sliding windows and across contig 11 for a) the resistant pool and b) the susceptible pool. The black lines indicate the delineations of the gap region, while the orange lines indicate the position of the Pasteuria Resistance Complex.

**Figure S3**: Number of SNPs called in 50kb sliding windows and across contig 11 for a) the resistant pool and b) the susceptible pool. The black lines indicate the delineations of the gap region, while the orange lines indicate the position of the Pasteuria Resistance Complex.

**Figure S4:** Manhattan plots representing the -log10(p-values) of the Fisher Exact Test conducted on allele frequencies between the resistant and susceptible pool as a function of genomic position across contig 11 for TNRA21 offspring depending on different filtering strategies based on allelic frequencies: a) Showing no filtering on the SNPs; b) SNPs filtered so their frequencies in the susceptible pools are over 90% (i.e., susceptible pool almost homozygote), and c) SNPs filtered so their frequencies in the resistant pools are over 90% (i.e., resistant pool almost homozygote). The orange lines indicate the coordinates of the Pasteuria Resistance Complex. SNPs in green are those retained for the investigation of the candidate gene for the G locus.

**Figure S5:** Manhattan plots representing the -log10(p-values) of the Fisher Exact Test conducted on alleles frequencies between the resistant and susceptible pool as a function of genomic position across the whole genome for each triplicate of TNRA21 offspring.

**Table S1:** Metrics generated by the co-occurrence test on attachment patterns between the *P. ramosa* isolate C1, C19 and P15 on a panel of 174 genotypes of *D. magna.* For each pair of *P. ramosa* isolates we show the number of attachments for the first isolate, the number of attachments for the second isolate, the number of observed co-occurrences (between *P; ramosa* strains), the number of expected co-occurrences, the probability of co-occurrence, the p-value associated with a negative co-occurrence (p-value -), the p-value associated with a positive co-occurrence (p-value +), the corrected p-value associated with a negative co-occurrence (FDR -), and the corrected p-value associated with a positive co-occurrence (FDR +). P-values were corrected using Benjamini & Hochberg false discovery rate (Benjamini and Hochberg 1995). Bold values refer to significant associations.

**Table S2:** Presentation of the 39 *Daphnia magna* clones used in the breeding experiment along with their resistotype for the attachment of *P. ramosa* isolates C1, C19, P15 hindgut (P15H) and P15 foregut (P15F). R stands for resistant and S for susceptible. The last two columns indicate if sexually produced F1 offspring were produced by selfing the clone.

**Table S3:** Percentage of observed and expected P15 foregut attachments (P15F) based on gene model 1 or gene model 2 for the G locus in the breeding experience. Clones that did not show segregation of resistance for P15 foregut attachment and are a) homozygous recessive at the C-locus (cc) and b) homozygous dominant (CC) or heterozygous (Cc) at the C-locus and homozygous at the G-locus (GG or gg depending on what model is considered). Clones that show segregation of resistance to P15 foregut attachment and are c) heterozygous (Cc) at the C-locus and homozygous recessive (gg) at the G-locus and d) homozygous dominant (CC) at the C-locus and heterozygous (Gg) at the G-locus. Some clones produced no selfed offspring, but showed P15 foregut attachment (clones FI-GE-3-11, PL-KNP-P4, PL-W2-1, US-SP131-1, US-SP15-1, US-SP163-1, US-SP221-1 and US-SP6-13).

**Table S4:** Pool description in terms of parental origin, resistotype for P15 foregut attachment, pool size and number of sequences

## References

Almeida R, Amado M, David L, Levery SB, Holmes EH, Merkx G, Van Kessel AG, Rygaard E, Hassan H, Bennett E, et al. 1997. A Family of Human β4-Galactosyltransferases. Journal of Biological Chemistry 272:31979–31991.

Ameline C, Bourgeois Y, Vögtli F, Savola E, Andras J, Engelstädter J, Ebert D. 2021. A Two-Locus System with Strong Epistasis Underlies Rapid Parasite-Mediated Evolution of Host Resistance. Molecular Biology and Evolution 38:1512–1528.

Audfray A, Varrot A, Imberty A. 2013. Bacteria love our sugars: Interaction between soluble lectins and human fucosylated glycans, structures, thermodynamics and design of competing glycocompounds. Comptes Rendus Chimie 16:482–490.

Benjamini Y, Hochberg Y. 1995. Controlling the False Discovery Rate: A Practical and Powerful Approach to Multiple Testing. Journal of the Royal Statistical Society Series B: Statistical Methodology 57:289–300.

Bento G, Fields PD, Duneau D, Ebert D. 2020. An alternative route of bacterial infection associated with a novel resistance locus in the Daphnia–Pasteuria host–parasite system. Heredity 125:173–183.

Bento G, Routtu J, Fields PD, Bourgeois Y, Du Pasquier L, Ebert D. 2017. The genetic basis of resistance and matching-allele interactions of a host-parasite system: The *Daphnia magna-Pasteuria ramosa* model. PLoS Genet 13:e1006596.

Beraldi D, McRae AF, Gratten J, Pilkington JG, Slate J, Visscher PM, Pemberton JM. 2007. Quantitative trait loci (QTL) mapping of resistance to strongyles and coccidia in the free-living Soay sheep (*Ovis aries*). International Journal for Parasitology 37:121–129.

Bever JD, Mangan SA, Alexander HM. 2015. Maintenance of Plant Species Diversity by Pathogens. Annu. Rev. Ecol. Evol. Syst. 46:305–325.

Bourgeois Y, Fields PD, Bento G, Ebert D. 2021. Balancing Selection for Pathogen Resistance Reveals an Intercontinental Signature of Red Queen Coevolution. Molecular Biology and Evolution 38:4918–4933.

Carius HJ, Little TJ, Ebert D. 2001. Genetic variation in a host-parasite association: potential for coevolution and frequency-dependent selection. Evolution 55:1136–1145.

Cornetti L, Fields PD, Du Pasquier L, Ebert D. 2024. Long-term balancing selection for pathogen resistance maintains trans-species polymorphisms in a planktonic crustacean. Nat Commun 15:5333.

De Vries T, Knegtel RMA, Holmes EH, Macher BA. 2001. Fucosyltransferases: structure/function studies. Glycobiology 11:119R–128R.

Dexter E, Fields P, Ebert D. 2023. Uncovering the Genomic Basis of Infection Through Co-genomic Sequencing of Hosts and Parasites. Molecular Biology and Evolution 40:msad145.

Dodds PN, Lawrence GJ, Catanzariti A-M, Teh T, Wang C-IA, Ayliffe MA, Kobe B, Ellis JG. 2006. Direct protein interaction underlies gene-for-gene specificity and coevolution of the flax resistance genes and flax rust avirulence genes. Proc. Natl. Acad. Sci. U.S.A. 103:8888–8893.

Duneau D, Luijckx P, Ben-Ami F, Laforsch C, Ebert D. 2011. Resolving the infection process reveals striking differences in the contribution of environment, genetics and phylogeny to host-parasite interactions. BMC Biol 9:11.

Ebert D. 2022. Daphnia as a versatile model system in ecology and evolution. EvoDevo 13:16.

Ebert D, Fields PD. 2020. Host–parasite co-evolution and its genomic signature. Nat Rev Genet 21:754– 768.

Eleftherianos I, Tafesh-Edwards G, Mohamed A. 2022. Pathogen infection routes and host innate immunity: Lessons from insects. Immunology Letters 247:46–51.

Fenton A, Antonovics J, Brockhurst MA. 2012. Two-step infection processes can lead to coevolution between functionally independent infection and resistance pathways: coevolution under multistep infection processes. Evolution 66:2030–2041.

Feron R. 2020. RomainFeron/PSASS: Psass 3.0.1. Available from: https://zenodo.org/record/3702337

Fredericksen M, Ameline C, Krebs M, Hüssy B, Fields PD, Andras JP, Ebert D. 2021. Infection phenotypes of a coevolving parasite are highly diverse, structured, and specific. Evolution 75:2540–2554.

Fredericksen M, Fields PD, Du Pasquier L, Ricci V, Ebert D. 2023. QTL study reveals candidate genes underlying host resistance in a Red Queen model system. PLoS Genet 19:e1010570.

Griffith DM, Veech JA, Marsh CJ. 2016. cooccur : Probabilistic Species Co-Occurrence Analysis in *R*. J. Stat. Soft. [Internet] 69. Available from: http://www.jstatsoft.org/v69/c02/

Gu Z, Gu MZ, GlobalOptions I. 2022. Package ‘circlize.’

Hall MD, Bento G, Ebert D. 2017. The Evolutionary Consequences of Stepwise Infection Processes. Trends in Ecology & Evolution 32:612–623.

Hamilton WD. 1982. Pathogens as Causes of Genetic Diversity in their Host Populations. In: Anderson RM, May RM, editors. Population Biology of Infectious Diseases. Berlin, Heidelberg: Springer Berlin Heidelberg. p. 269–296. Available from: http://link.springer.com/10.1007/978-3-642-68635-1_14

Hite JL. 2020. Host age alters disease life history. A case study in zooplankton and a castrating pathogen. Functional Ecology 34:1522–1524.

Huessy B, Bumann D, Ebert D. 2024. Ectopical expression of bacterial collagen-like protein supports its role as adhesin in host–parasite coevolution. R. Soc. Open Sci. 11:231441.

Iyer J, Grüner AC, Rénia L, Snounou G, Preiser PR. 2007. Invasion of host cells by malaria parasites: a tale of two protein families. Molecular Microbiology 65:231–249.

Izhar R, Gilboa C, Ben-Ami F. 2020. Disentangling the steps of the infection process responsible for juvenile disease susceptibility. Functional Ecology 34:1551–1563.

Kofler R, Pandey RV, Schlötterer C. 2011. PoPoolation2: identifying differentiation between populations using sequencing of pooled DNA samples (Pool-Seq). Bioinformatics 27:3435– 3436.

Kurtz J, Schulenburg H, Reusch TBH. 2016. Host–parasite coevolution—rapid reciprocal adaptation and its genetic basis. Zoology 119:241–243.

Li H. 2013. Aligning sequence reads, clone sequences and assembly contigs with BWA-MEM. Available from: http://arxiv.org/abs/1303.3997

Li J, Hsu H-C, Mountz JD, Allen JG. 2018. Unmasking Fucosylation: from Cell Adhesion to Immune System Regulation and Diseases. Cell Chemical Biology 25:499–512.

Lievens EJP, Perreau J, Agnew P, Michalakis Y, Lenormand T. 2018. Decomposing parasite fitness reveals the basis of specialization in a two-host, two-parasite system. Evolution Letters 2:390– 405.

Loot G, Poulet N, Reyjol Y, Blanchet S, Lek S. 2004. The effects of the ectoparasite *Tracheliastes polycolpus* (Copepoda: Lernaeopodidae) on the fins of rostrum dace (*Leuciscus leuciscus burdigalensis*). Parasitology Research 94:16–23.

Luijckx P, Fienberg H, Duneau D, Ebert D. 2013. A Matching-Allele Model Explains Host Resistance to Parasites. Current Biology 23:1085–1088.

Lydecker HW, Etheridge B, Price C, Banks PB, Hochuli DF. 2019. Landscapes within landscapes: A parasite utilizes different ecological niches on the host landscapes of two host species. Acta Tropica [Internet]. Available from: https://linkinghub.elsevier.com/retrieve/pii/S0001706X18315572

Ma B, Simala-Grant JL, Taylor DE. 2006. Fucosylation in prokaryotes and eukaryotes. Glycobiology 16:158R–184R.

Martins NE, Faria VG, Teixeira L, Magalhães S, Sucena É. 2013. Host Adaptation Is Contingent upon the Infection Route Taken by Pathogens. Schneider DS, editor. PLoS Pathog 9:e1003601.

Metzger CMJA, Luijckx P, Bento G, Mariadassou M, Ebert D. 2016. The Red Queen lives: Epistasis between linked resistance loci: BRIEF COMMUNICATION. Evolution 70:480–487.

Mouritsen KN, Poulin R. 2005. Parasites boosts biodiversity and changes animal community structure by trait-mediated indirect effects. Oikos 108:344–350.

Pigeault R, Isaïa J, Yerbanga RS, Dabiré KR, Ouédraogo J-B, Cohuet A, Lefèvre T, Christe P. 2020. Different distribution of malaria parasite in left and right extremities of vertebrate hosts translates into differences in parasite transmission. Sci Rep 10:10183.

Råberg L. 2023. Human and pathogen genotype-by-genotype interactions in the light of coevolution theory.Gibson G, editor. PLoS Genet 19:e1010685.

Rini JM, Moremen KW, Davis BG, Esko JD. 2022. Glycosyltransferases and Glycan-Processing Enzymes. In: Essentials of Glycobiology [Internet]. 4th edition. Cold Spring Harbor Laboratory Press.

Robinson JT. 2011. Integrative genomics viewer. correspondence 29.

Routtu J, Ebert D. 2015. Genetic architecture of resistance in Daphnia hosts against two species of host-specific parasites. Heredity 114:241–248.

Santos JL, Ebert D. 2023. The limits of stress-tolerance for zooplankton resting stages in freshwater ponds. Oecologia 203:453–465.

Schlötterer C, Tobler R, Kofler R, Nolte V. 2014. Sequencing pools of individuals — mining genome-wide polymorphism data without big funding. Nat Rev Genet 15:749–763.

Shea JE, Hensel M, Gleeson C, Holden DW. 1996. Identification of a virulence locus encoding a second type III secretion system in Salmonella typhimurium. Proc. Natl. Acad. Sci. U.S.A. 93:2593– 2597.

Sherman RM, Salzberg SL. 2020. Pan-genomics in the human genome era. Nat Rev Genet 21:243–254.

Spurgin LG, Richardson DS. 2010. How pathogens drive genetic diversity: MHC, mechanisms and misunderstandings. Proc. R. Soc. B. 277:979–988.

Su C, Howe DK, Dubey JP, Ajioka JW, Sibley LD. 2002. Identification of quantitative trait loci controlling acute virulence in *Toxoplasma gondii*. Proc. Natl. Acad. Sci. U.S.A. 99:10753–10758.

Turner AK, Begon M, Jackson JA, Bradley JE, Paterson S. 2011. Genetic Diversity in Cytokines Associated with Immune Variation and Resistance to Multiple Pathogens in a Natural Rodent Population. PLoS Genetics 7:e1002343.

Vasimuddin Md, Misra S, Li H, Aluru S. 2019. Efficient Architecture-Aware Acceleration of BWA-MEM for Multicore Systems. Available from: http://arxiv.org/abs/1907.12931

Webster JP, Woolhouse MEJ. 1998. Selection and strain specificity of compatibility between snail intermediate hosts and their parasitic schistosomes. Evolution 52:1627–1634.

Wen M, Zhang Y, Wang Siyu, Hu F, Tang Congjia, Li Q, Qin Q, Tao M, Zhang C, Zhao R, et al. 2022. Sex locus and sex markers identification using whole genome pool-sequencing approach in the largemouth bass (*Micropterus Salmoides L*.). Aquaculture 559:738375.

