## Supplementary figures and images for "Genetic basis of resistance in hosts facing alternative infection strategies by a virulent bacterial pathogen"

### Figure S1

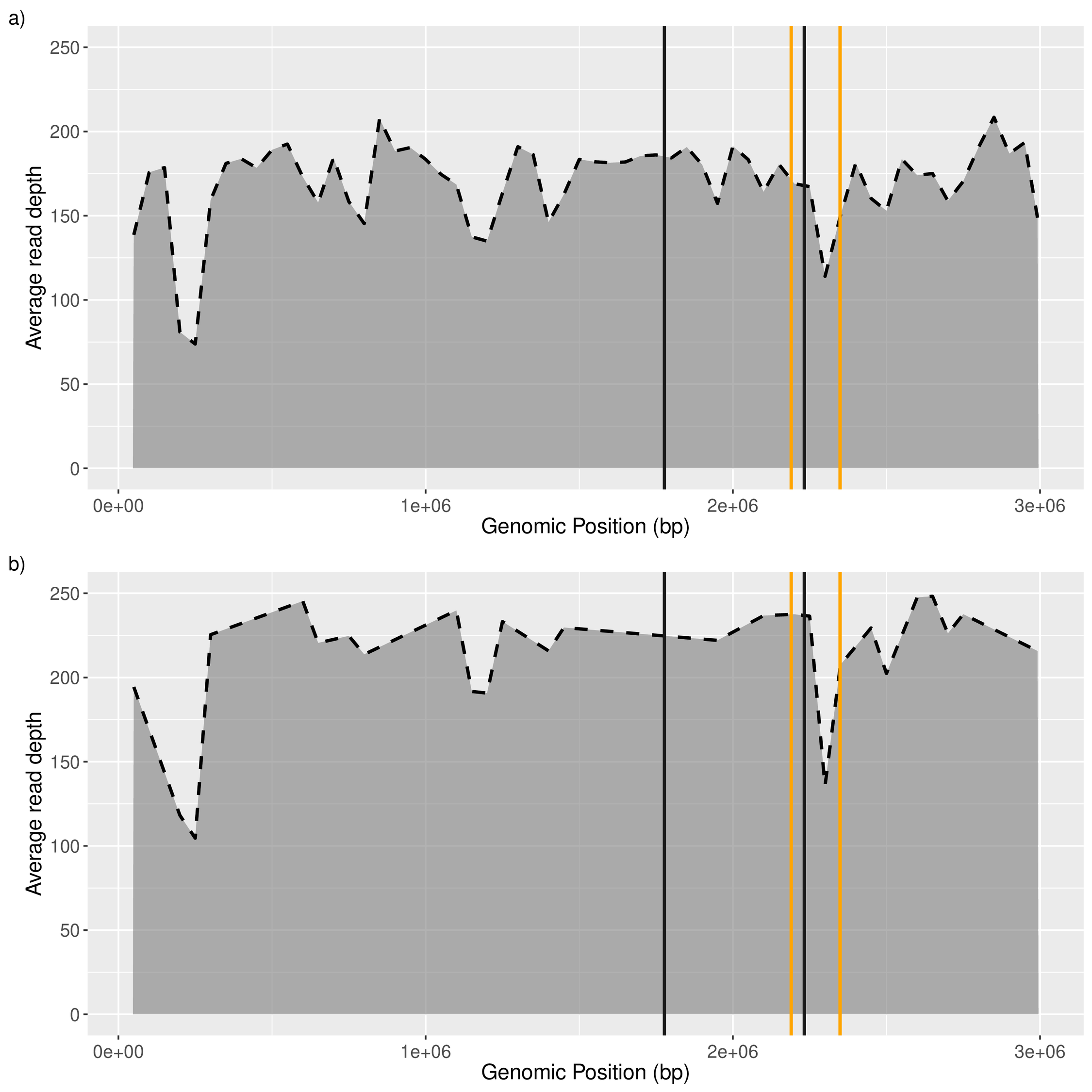

### Figure S2

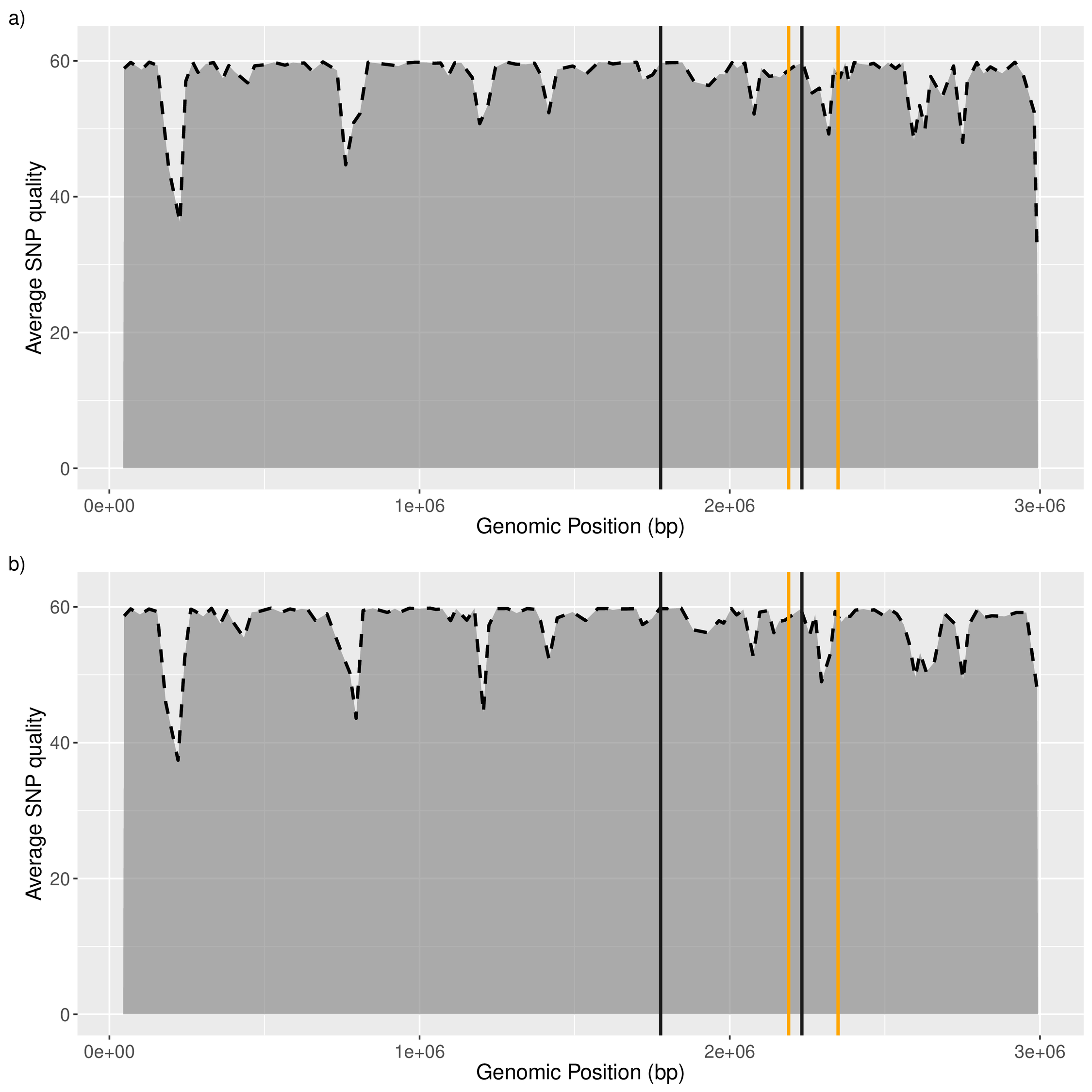

### Figure S3

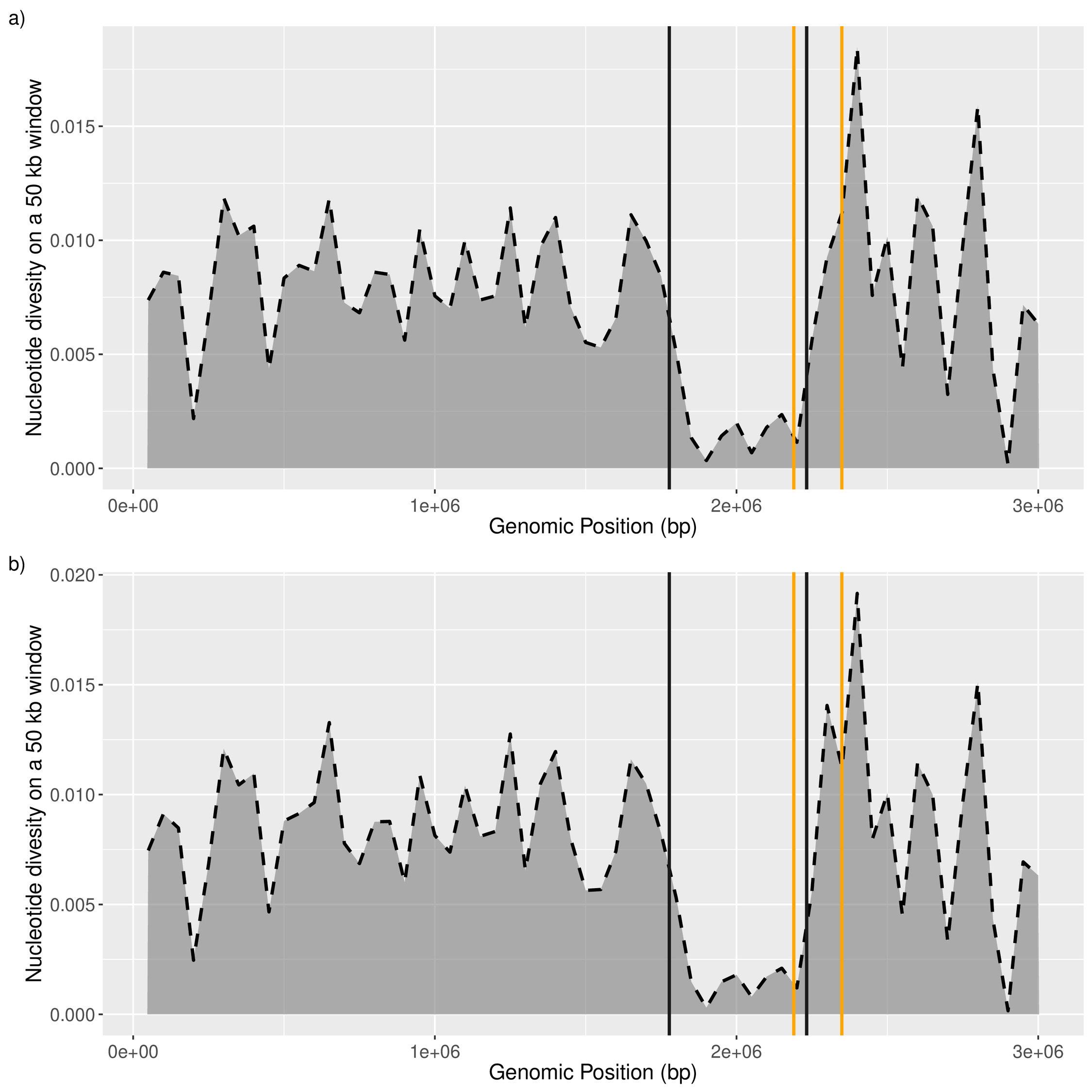

### Figure S4

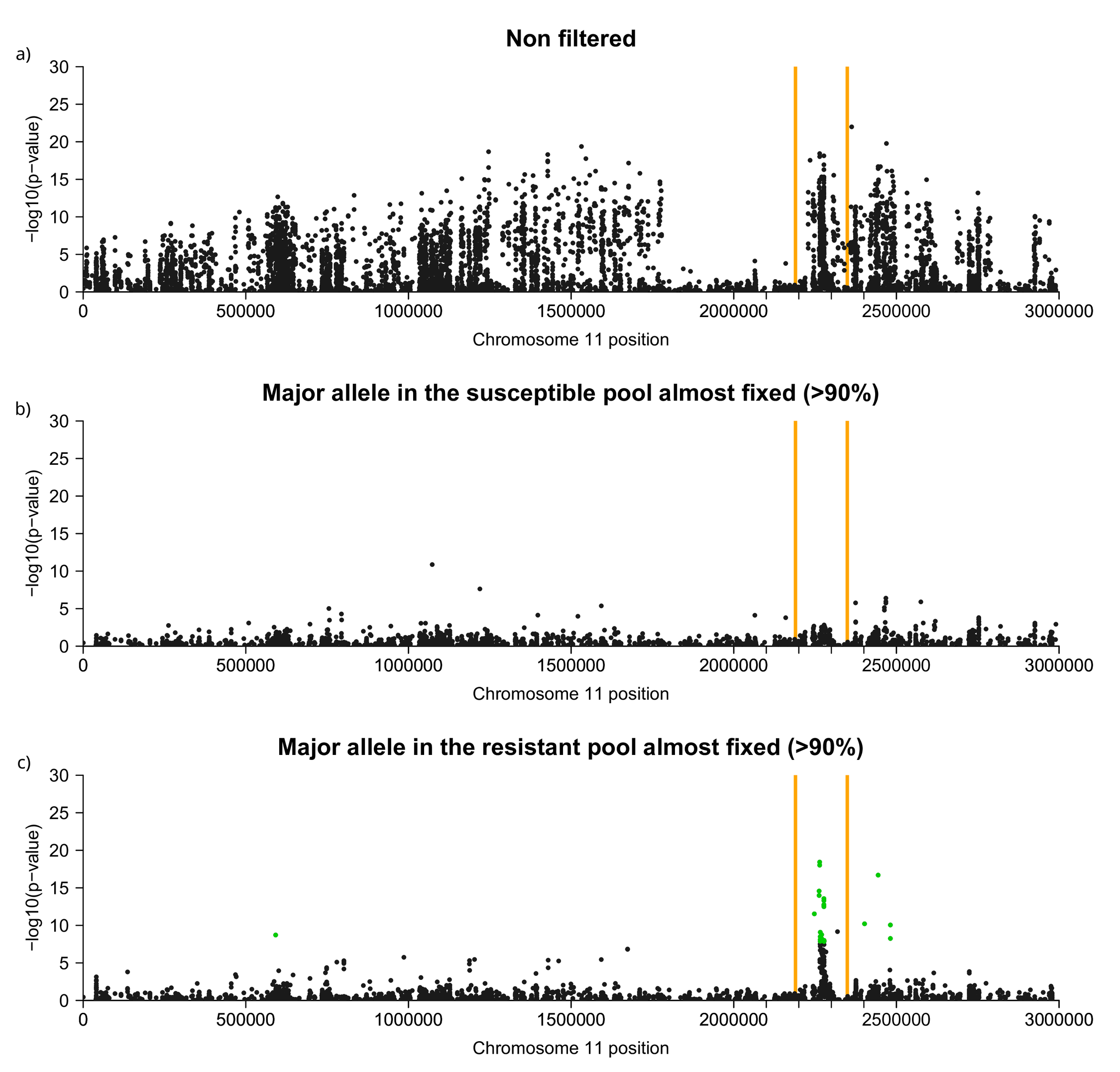

### Figure S5

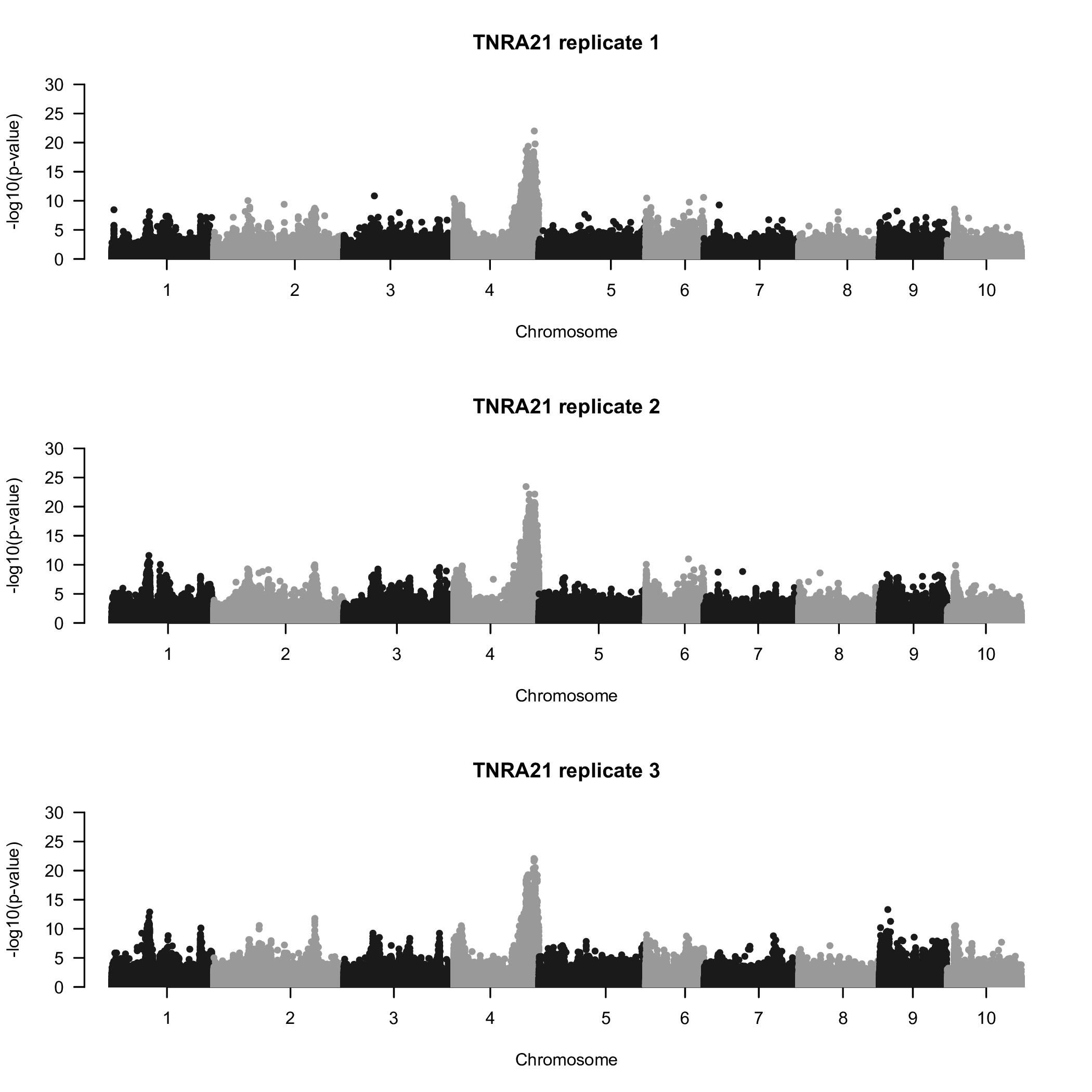
